# Stress promotes RNA G-quadruplex folding in human cells

**DOI:** 10.1101/2022.03.03.482884

**Authors:** Prakash Kharel, Marta Fay, Ekaterina V. Manasova, Paul J. Anderson, Aleksander V. Kurkin, Junjie U. Guo, Pavel Ivanov

**Affiliations:** Brigham and Women’s Hospital, Harvard Medical School, Boston, MA 02115, USA; Chemistry Department of Lomonosov Moscow State University, 119991, Moscow, Russia; Harvard Medical School Initiative for RNA Medicine, Boston, MA 02115, USA; Department of Neuroscience, Yale School of Medicine, New Haven, CT 06520, USA

## Abstract

Guanine-rich nucleic acids can fold into G-quadruplex (G4) structures. Although endogenous RNAs contain sequences that can fold into RNA G4s (rG4s) *in vitro*, their folding and functions *in vivo* are not well understood. We show that the folding of putative rG4s in human cells into bona fide rG4 structures is dynamically regulated by stress. By using a high-throughput approach based on differential reactivity of dimethyl sulfate (DMS) towards Gs within folded vs unfolded rG4s, we identified hundreds of endogenous rG4s whose folding is promoted by cellular stress and validated them using a newly developed rG4-specific ligand. Stress-dependent rG4s are enriched in mRNA 3′-untranslated regions, suggesting their role in regulating mRNA stability under stress. Lastly, rG4 folding is reversible upon stress removal or adaptation. Our study show that rG4s function as regulatory elements in regulating mRNA stability and cellular stress response.

**One-Sentence Summary:** RNA G-quadruplexes assemble under stress in human cells

## Main text

G-quadruplexes (G4s) are nucleic acid secondary structures formed by the stacking of two or more G-quartets, each formed by a square planar arrangement of four guanines (Gs) connected by H-bonding^1^ **(Fig. 1a)** and stabilized by the presence of monovalent cations such as K^+^ and Na^+2^. Potential G4-forming sequences are enriched in regulatory regions of both genome and transcriptome ^3,4^, and have been implicated in many cellular processes, and the number of biological functions attributed to them continues to grow^5^. Computational analyses and high throughput studies of the human transcriptome suggest that thousands of putative rG4s are found in mRNAs^6-8^ and they may play important roles in the regulation of mRNA maturation, transport, and translation^5,9,10^, and are associated with human diseases^10^. Potential rG4 sequences are more prevalent in the 5′ and 3′ untranslated regions (UTR) of mRNAs-partly due to the strong depletion of such sequences in the coding sequence (CDS). rG4s in 5′ UTRs are associated with modulation of translation^11-13^, while rG4s within CDS have been shown to impede translation and contribute to mRNA decay^14^. On the other hand, rG4 sequences in the 3′ UTR influence microRNA targeting^15^, RNA localization^16^, and alternative polyadenylation^12^. Additionally, recent studies suggest that RNA secondary structures can act as molecular seeds to assemble RBPs that can eventually create granular bodies^17^. Collectively, these findings indicate that rG4 sequences perform crucial regulatory functions during posttranscriptional events and RNA metabolism in eukaryotes.

**Figure 1.**
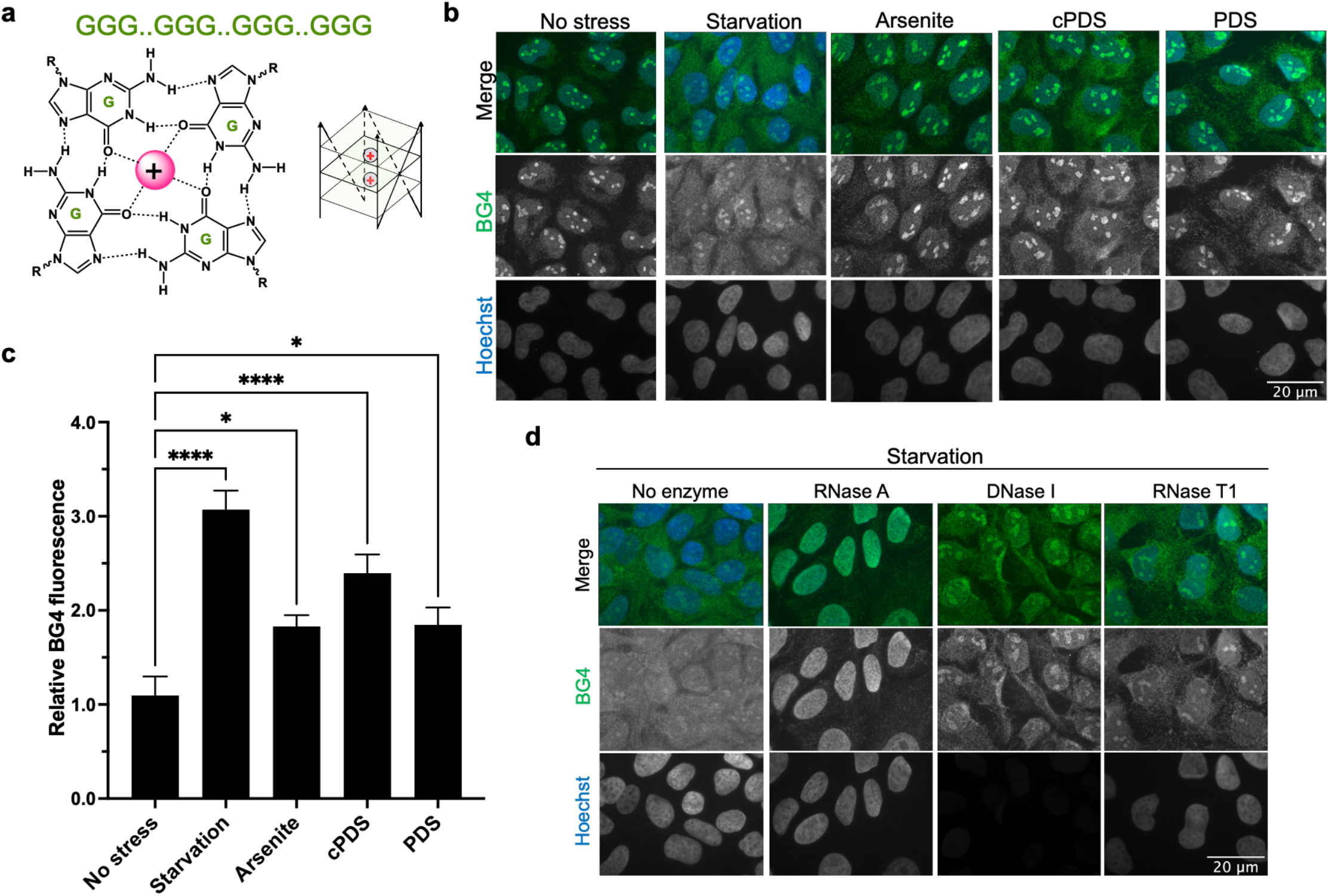
RNA G-quadruplex formation is enhanced under stress in cells. **a**, A square planar G-quartet arrangement stabilized by a monovalent cation (H-bonds and cation-oxygen coordination is shown with dotted bonds), and a schematic of 3-tier G-quadruplex structure stabilized by monovalent cations. The schematic depicts the stacking of 3 G-quartets (light green planes) with RNA-backbone in a parallel orientation. **b**, BG4 staining in U2OS cells demonstrate both RNA and DNA can form G4s in U2OS cells and rG4 folding (cytosolic BG4 fluorescence) in the U2OS cells is increased under various stress conditions as indicated by a more intense fluorescence signals of BG4 antibody under 100 µM sodium arsenite mediated oxidative stress and starvation stress, rG4 and G4 stabilizing ligands carboxypyridostatin (cPDS) and pyridostatin (PDS) were used as positive controls. **c**, Quantification of fluorescence intensity of BG4 signals in 100 cells per condition of 3 independent experiments, and **d**, RNase A treatment almost completely abolished the cytoplasmic BG4 signaling, while DNase I treatment minimized nuclear BG4 staining, and RNase T1 treatment had little effect on resultant cytosolic BG4 fluorescence clearly suggesting cellular RNA is folded in rG4 under stress.

Despite the accumulating evidence supporting a regulatory role for rG4 sequences, whether they act as single-stranded RNA motifs or as folded rG4 structures has been unclear. Efforts have been made to detect rG4 folding in cells using different approaches, have led to distinct conclusions. On one hand, G4-specific antibodies^18^ and rG4-specific chemical probes^19,20^ have detected folded rG4 structures in human cells^18,21^ and in plants^22^. These reagents, however, could potentially introduce biases to the rG4 folding landscape by inducing or stabilizing rG4 formation. Additionally, the possibility of rG4 folding during the process of cell fixation and visualization cannot be completely ruled out. Furthermore, rG4 antibodies such as BG4, which has been widely used for detecting rG4s in cells^21^, have been shown to cross-react with non-G4 sequences^23^. Thus, these studies are unable to determine the fraction of rG4 sequences that fold into rG4 structures. Methods based on RNA chemical structure probing present a more quantitative landscape of rG4 folding in living cells. One method is based upon the ability of 2-methylnicotinic acid imidazolide (NAI) to methylate the 2′-hydroxyl of terminal G residues within G-tracts when rG4 is in a folded state. These modifications can be quantified by analyzing the reverse transcription stops^22,24^. A second method uses dimethyl sulfate (DMS) to methylate the N7 position of guanine residues (N7G) that are not part of rG4 structures^24^(**Fig. 1a**). Once methylated at N7, these G nucleotides cannot fold back to rG4s *in vitro*, thereby alleviating RT stalling by folded rG4 structures. Both methods have previously been applied to the potential rG4s in the poly(A)^+^ transcriptome. The results have shown that although up to 50% of mammalian mRNAs can readily assemble rG4s *in vitro*, they are overwhelmingly unfolded in yeast and mammalian cells^*24*^. While DMS and SHAPE reagents, especially at high concentrations, can also potentially influence rG4 folding in cells, a recent study has shown folded rG4s in *Arabidopsis* by using a similar SHAPE-based approach^22^, indicating that chemical probing itself does not preclude the detection of rG4s *in vivo*.

The prevalence of mammalian rG4s and their unfolded state under normal conditions raise the possibility that their folding may be regulated by cellular and/or environmental signals. In this report, we found the rG4 folding is modulated by stress in human cells. Using chemical structure probing in combination with high-throughput sequencing and RT stall analysis, we identified rG4s that have a propensity to fold into rG4s under different stress environments. Next, we developed a small-molecule probe that can selectively pull down rG4s from the cellular transcriptome and validated the identity of bona-fide rG4s found by high-throughput sequencing. Interestingly, stress-induced global rG4 folding is reversible upon removal of stress suggesting that rG4s are dynamic stress-responsive elements. Our findings provide the first direct evidence of stress-dependent global folding of rG4s in human cells, introducing rG4 motifs as important players in the regulation of cellular processes under stress.

### Elevated rG4 folding in human cells under stress

To test whether rG4 folding may be regulated by cellular states, human osteosarcoma U2OS cells were fixed and probed with G4-specfic antibody (BG4)^21^, which showed predominantly nuclear but some cytoplasmic signals (**Fig. 1b**, panel 1). Interestingly, U2OS cells cultured in starvation media (Hank’s balanced salt solution, HBSS media) for two hours before fixation showed significantly increased BG4 signal intensity in the cytoplasm (**Fig. 1b-c**), suggesting that rG4s become more folded upon starvation. To test whether other cellular stress may also induce rG4 folding, we induced oxidative stress in U2OS cells by treating them with sodium arsenite for one hour, and observed a ∼2-fold increase in BG4 signal intensity (**Fig. 1b-c**). As validations for the specificity of BG4, known rG4-stabilizing ligands carboxypyrodistatin (cPDS) and pyridostatin (PDS) also enhanced BG4 staining **(Fig. 1b-c**). While DNase I treatment reduced BG4 signal in the nucleus, cytoplasmic BG4 signal was resistant to DNase I but was instead sensitive to RNase A treatment (**Fig. 1d**). In contrast, RNase T1, which targets guanosines in unfolded RNA regions, has a minimal effect in reducing BG4 signals, strongly indicating that increased cytoplasmic BG4 signal under stress was indeed due to rG4 folding. Quantitatively similar results were observed using transformed African green monkey kidney cells (COS7) (**fig. S1**).

### Poly-A RNAs demonstrate enrichment of *in vivo* rG4 folding in mRNA 3’ UTRs under stress

To systematically investigate the endogenous rG4s that are folded in U2OS cells under stress, we first used a reverse transcriptase (RT)-stop profiling approach to identify endogenous rG4 regions in the transcriptome^24^. During primer extension, rG4s cause RT stalling in a K^+^-dependent manner, generating truncated cDNA fragments that can be sequenced in a high-throughput manner. As expected, strong RT stops were highly enriched (70%) at G nucleotides (**Fig. 2b**), indicating that rG4s present the primary barrier for RT. Consistent with the stabilization of rG4 structures by K^+^, the fraction of stops at G nucleotides was even higher (86%) among the 2,224 RT stops that were ≥2-fold stronger in K+ compared to Na^+^. In agreement with previous studies^7^, rG4 regions were significantly enriched in 3′ UTRs, with 72% rG4 RT stops compared to 42% of all RT stops residing in 3′ UTRs (**Fig. 2c**).

**Figure 2.**
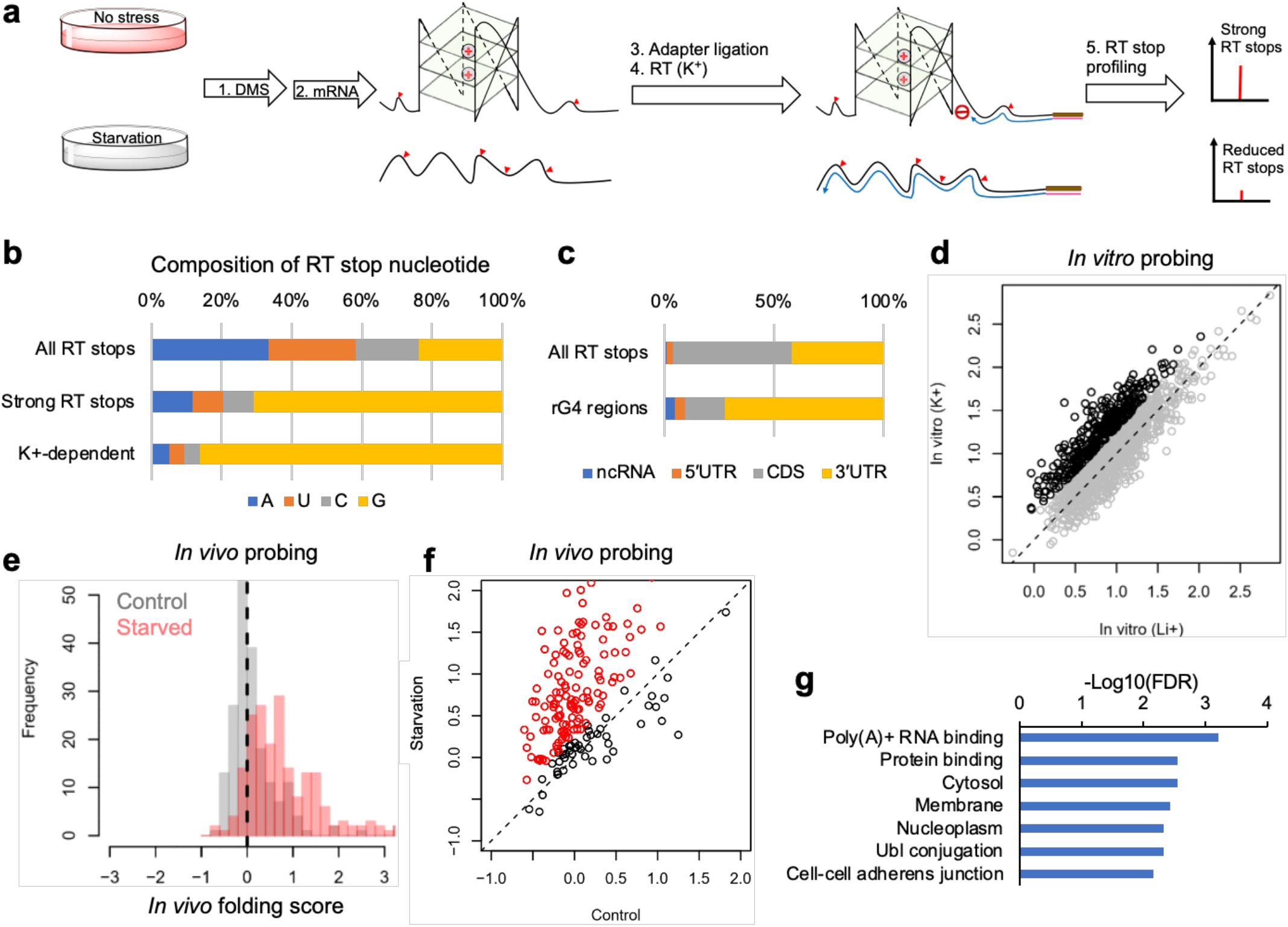
DMS mediated unbiased structure probing shows mRNA rG4s are folded under stress in vivo. **a**, A schematic of DMS-seq and RT-stop profiling to identify the bona-fide rG4s in U2OS cells under starvation stress. **b**, RT stops are highly enriched in G residues. **c**, RT stops are selectively enriched in mRNA 3’UTRs. **d**, On average, RT stops in 150 mM K+ folded followed by DMS modification mRNA are 1.6-fold stronger than that in 150 mM Li+ folded followed by DMS modified mRNA. **e-f**, in vivo rG4 folding increases significantly in starved U2OS cells but not in the unstressed cells. **g**, mRNAs containing starvation-induced folded rG4s are enriched in those encoding RNA-binding proteins, ubiquitin ligases, and both cytosolic and membrane proteins.

Next, we quantified the folding states of these endogenous rG4 regions *in vitro* and *in vivo* using DMS probing. DMS methylates the N7 positions of G nucleotides that are accessible and not engaged in G-quartets thereby reducing RT stalling during primer extension. In contrast, the N7 positions in G-quartets are protected from DMS (**Fig. 2a**). Indeed, polyA-selected mRNAs folded and DMS-modified in the presence of 150 mM K^+^ caused on average 1.6-fold stronger RT stops than those folded and DMS-modified in the presence of 150 mM Li^+^ (**Fig. 2d**). 335 out of 1,346 (25%) rG4 regions showed ≥2 fold difference in DMS accessibility between K^+^ and Li^+^ conditions (**Fig. 2d**, black), which we carried forward for *in vivo* DMS probing analysis.

To quantify rG4 folding in vivo, we performed DMS probing in U2OS cells under either basal or starvation conditions and calculated the *in vivo* folding scores for each of 196 rG4 regions with sequencing coverage under all conditions. Consistent with previous results^24^, the vast majority of rG4 regions in unstressed cells *in vivo* were highly accessible to DMS modifications, with a median folding score of -0.01, indicating that they were in a predominantly unfolded state similar to the *in vitro* state without K+. Strikingly, we observed strong protection of these rG4 regions from DMS modifications after starvation, resulting in a median *in vivo* folding score of 0.64, suggesting that many rG4s became partly if not completely folded (**Fig. 2e**). Starvation-induced rG4 folding appeared to be widespread, with 72% rG4s showing an increase of *in vivo* folding scores of ≥0.25 (**Fig. 2f**). mRNAs containing starvation-induced folded rG4s were enriched in those encoding RNA-binding proteins, ubiquitin ligases, and both cytosolic and membrane proteins (**Fig. 2g**). Collectively, our unbiased DMS-seq analysis showed that starvation induced widespread and stable folding of rG4 structures, raising the possibility that they may regulate mRNA metabolism in a stress-dependent manner.

### *In vitro* validation of rG4 formation of selected mRNA oligos

To verify the capability of DMS-seq identified mRNA 3′ UTR G-rich regions to fold into rG4s, we analyzed the CD characteristics of respective RNA oligos under rG4 permissive and non-permissive conditions. All the tested candidates showed an expected CD characteristic of rG4s, indicating these sequences can indeed fold into rG4 (**fig. S3**). In addition, we used electromobility gel shift assay and verified their G4 forming ability by using a G4 specific dye. N-methyl mesoporphyrin IX (NMM) selectivity to bind parallel G4s was used to detect rG4s in acrylamide gels. Surprisingly, NMM stained some pG4-oligo bands both in non-denaturing K^+^ and Li^+^ folding conditions as well as under denaturing conditions, clearly indicating the stable nature of these rG4s (**fig. S4 and Fig. 3b**). Additionally, an NMM-fluorescence assay shows enhanced fluorescence of rG4-NMM complexes (**Fig. S5**). NMM fluorescence data support the CD and NMM gel staining results indicating the superstable nature of candidate rG4s.

**Figure 3.**
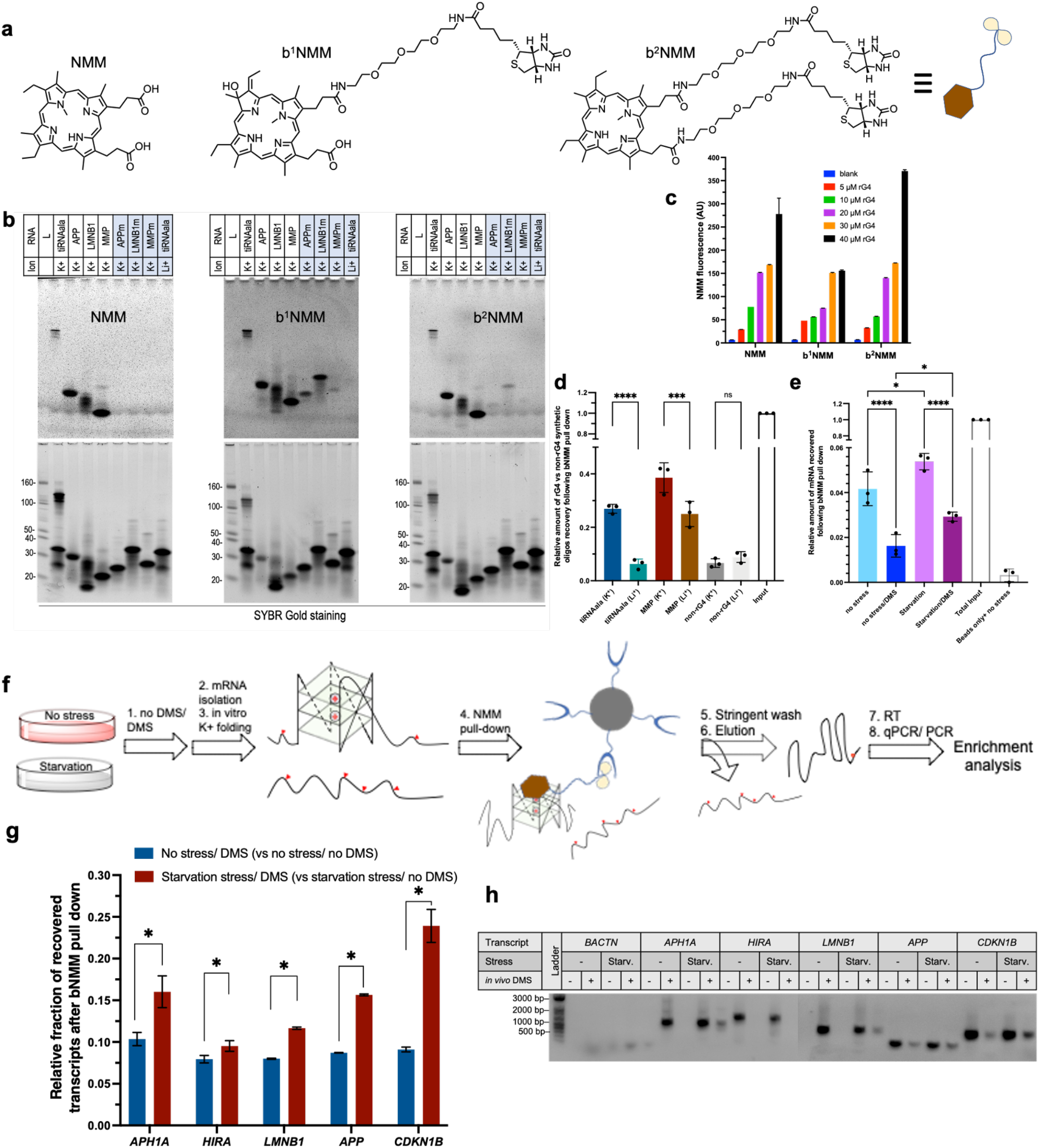
Biotin functionalized NMM (bNMM) acts a selective probe to pull down RNA G-quadruplexes. **a**, Chemical structure of NMM, a parallel G4 specific porphyrin (right) and biotin functionalized NMMs (synthesis and characterization in SI). **b**, gel staining demonstrate double biotinylated NMM shows similar rG4 staining behavior as its parent compound while single biotinylated version is less selective. **c**, Biotin functionalized NMMs show similar G4 affinity as their parent compound in NMM fluorescence assay, 5’tiRNA^Ala^ G4 is used for the selectivity test since 5’tiRNA^Ala^ -NMM characterization is already established. **d**, Validation of bNMM based rG4 pull-down approach using synthetic rG4 oligos. **e**, Validation of bNMM-rG4 pull-down approach using cellular mRNA. This also affirms that under starvation stress, more Gs are protected from DMS reactivity. **f**, Schematic of cellular rG4 enrichment and their quantification. Cellular mRNA (with or without the starvation stress and with (D) and without (-) the DMS reaction) was isolated and folded in presence of K^+^ before mixing with bNMM beads. The captured mRNA pool was eluted, reverse transcribed, and the enrichment was quantified using qPCR and PCR. **g**, Candidate mRNAs from DMSeq experiments show a higher abundance in the mRNA isolated from DMS treated stressed cells, clearly indicating their folding into rG4s *in vivo*. A higher CT value corresponds to the lower abundance of a particular transcript in the eluant (Raw CT values in supplementary figure). **h**, RT-PCR amplicons are run in 1% agarose gel to visualize the difference in the bNMM bound mRNA under similar conditions as in **g**.

### Biotinylated N-methyl mesoporphyrin (bNMM) as a tool to capture cellular rG4s

To validate DMS-seq-identified folded rG4 candidates, we developed a probe that can capture bona-fide rG4s from cell lysates. Because NMM preferentially binds to parallel G4s, we reasoned that biotin modification of NMM could provide a convenient tool to selectively pull down the folded rG4 from the cellular mRNA pool. A similar approach has been used to selectively pull down parallel dG4s (REF). In combination with DMS modification, biotinylated NMM provides a powerful tool to delineate *in vivo* rG4 formation. Two biotinylated derivatives of NMM, namely single biotinylated NMM (b^1^NMM) and double biotinylated NMM (b^2^NMM) were synthesized (**Fig. 3a**) with 18-atom long linker(s) to provide enough space to interact with large RNA molecules.

To test whether biotin modification impacts the ability of NMM to recognize the rG4s, we first compared their fluorescence behavior when bound to rG4s. Because we previously studied NMM fluorescence with 5’tiRNA^Ala^ rG4s^25^, we used the same platform to evaluate the rG4 recognition ability of biotin-modified NMMs. b^2^NMM showed comparable fluorescence behavior with that of unmodified NMM (**Fig. 3b**), demonstrated a comparable selectivity in rG4 recognition for gel staining (**Fig. 3c**), and was chosen for further downstream applications. bNMM provides a specific and efficient platform to select rG4s using *in vitro* synthesized short rG4 forming oligos. Previously well characterized rG4s (tiRNA^Ala26^ and MMP^27^) and control oligos were folded under K^+^ or Li^+^ environment and allowed to equilibrate with streptavidin-agarose bound bNMM. After washing, the bound RNA was eluted and spectrophotometrically quantified. As presented in **Fig. 3d**, we observed significant enrichment of bNMM pulled-down oligos under G4 permissive conditions clearly showing the ability of streptavidin-bound bNMM beads to pull down rG4s.

This approach was used to capture total rG4 mRNAs from stressed and unstressed cells with or without prior DMS treatment. The spectrophotometric quantification of pulled down and eluted mRNAs demonstrated a pattern of recovery that corroborates the idea that cellular rG4s are more folded under starvation conditions (**Fig. 3e**). If an rG4 region is folded *in vivo*, the N7 position of constituent Gs would be protected from DMS, which in turn allows such sequence to fold back to rG4 under G4 permissive condition (150 mM K^+^) *in vitro*. Consequently, bNMM trapping of total mRNA isolated from unstressed cells with or without the DMS treatment (-/D and -/-) versus those from the stressed cells (H/D and H/-), and their subsequent elution and quantification provides a snapshot of relative rG4 folding of individual mRNA *in vivo*. As schematically represented in **Fig. 3f**, total mRNA was isolated from U2OS cells under various conditions and was folded in presence of 150 mM KCl. Folded total mRNA was incubated with streptavidin agarose-bound bNMM to capture rG4-containing mRNAs. Bound mRNA was eluted, and relative abundance of bNMM-bound mRNA was determined using reverse transcription quantitative polymerase chain reaction (RT-qPCR) (**Fig. 3g-h**). Both of these experiments showed that a higher proportion of bNMM-bound rG4 forming mRNAs were available under starvation situation.

### 3’UTR rG4s containing mRNAs are more stable under stress

Because 3′ UTR structural elements are known to regulate mRNA decay^28,29^, we hypothesized that the induced folding of 3′ UTR rG4s might regulate the stability of those transcripts under stress. As demonstrated in the scheme **(Fig. 4a)**, transcriptional incorporation of 4-thio-uracil (4TU) allows newly synthesized RNAs to be tagged with biotin, which allows the separation of new transcripts from the older decaying transcripts^30^. The relative abundance of transcripts present in total vs non-biotinylated (decaying) transcript pools quantifies the decay kinetics of corresponding transcripts without influence of stress on transcription. To determine the connection between rG4 formation and mRNA stability, we performed pulse chase experiments with 4TU in U2OS cells with or without starvation stress. Total mRNA isolated at each time point was biotinylated and separated into two pools using streptavidin affinity capture. We observed that *LMNB1* and *HIRA* mRNAs, two rG4-containing mRNAs identified from DMS-seq, had significantly longer half-lives under stress while the total mRNA level is not significantly different between no stress vs stress conditions **(Fig. 4b)**. In contrast, such a difference was not observed for 28S rRNA used as control. These results suggest that rG4 folding under stress stabilizes the respective transcripts.

**Figure 4.**
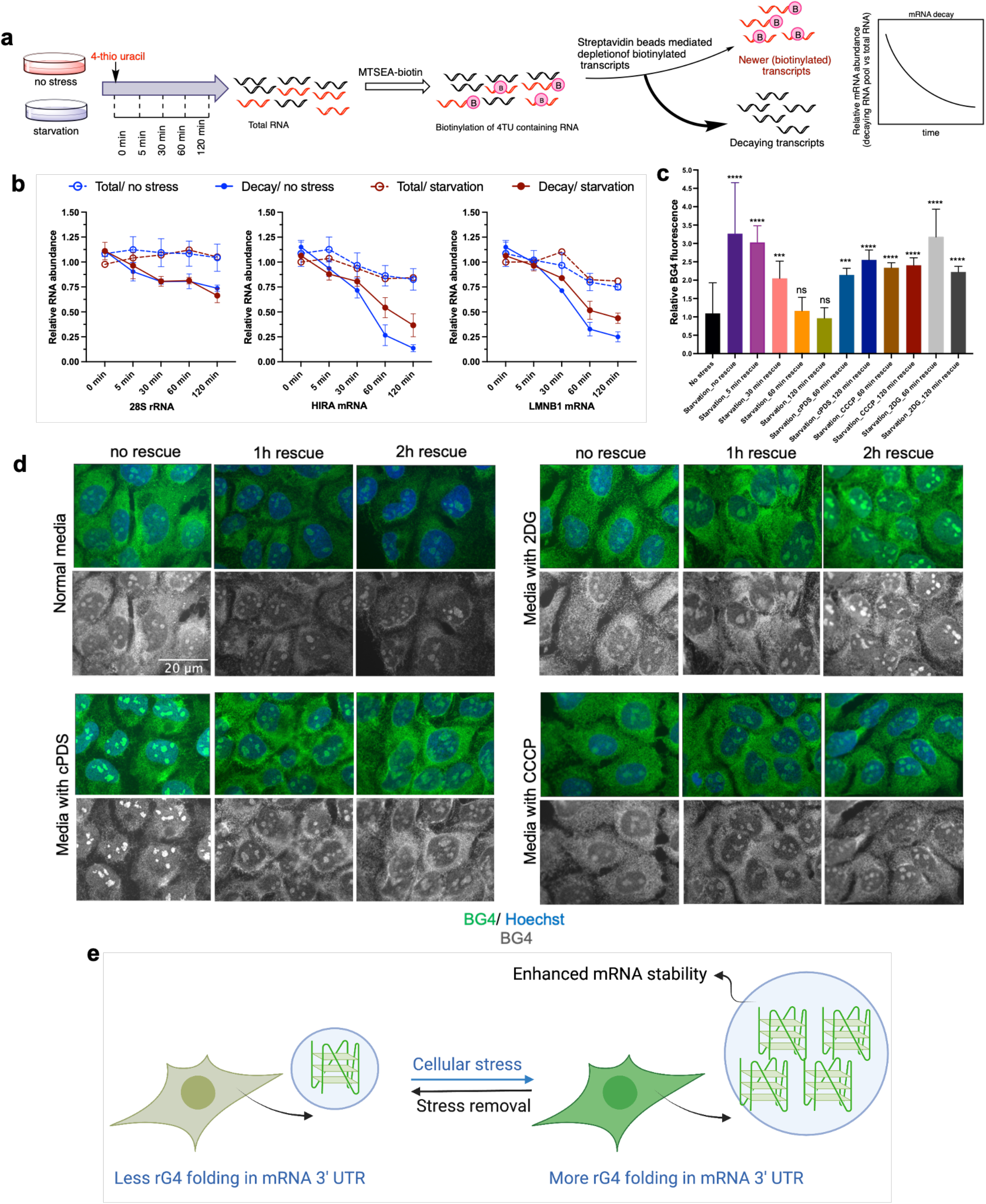
rG4s within the 3′ UTRs enhance mRNA stability under stress. **a**, Schematic of 4TU doping, MTSEA tagging, biotinylation, and decaying mRNA enrichment and analysis process. **b**, qPCR quantification of decaying mRNA (collected under indicated timepoint) shows candidate mRNAs (*HIRA* and *LMNB1*) are significantly more stable under starvation stress. **c**, Quantification of BG4 staining from starvation-rescue experiments (**Figure S6**). Starvation-rescue experiment demonstrates the reversibility of rG4 folding, and **d**, rG4 unfolding upon stress removal is largely paused by rG4 stabilizing cPDS, along with nutrient stresses created by 2DG (glycolysis inhibitor) and CCCP (oxidative phosphorylation inhibitor). Proposed model to demonstrate dynamic nature of stress responsive rG4 folding and their contribution in mRNA stability.

### rG4 folding under stress is reversible

Reversibility is one of the key features of cellular stress response mechanisms. Inability to reverse stress-induced changes upon stress resolution can perturb cellular physiology and contribute to disease. When folded *in vitro*, many rG4s represent unusually stable structures that are resistant to elevated temperatures, which begs the question whether *in vivo* folding of rG4s can be reversed when cells are no longer under stress. We observed that the elevated cytosolic rG4 signals were also decreased gradually, and eventually returned to a basal level within an hour of stress removal (**Fig. 4c-d, Fig S6**), indicating that the *in vivo* folding of rG4 is dynamic and reversibly regulated by cellular stress. To evaluate whether the unfolding dynamics of stress induced rG4s upon removal of stress can be modulated by using rG4 specific ligand, we treated the recovering cells with cPDS. Interestingly, the unfolding of rG4s was delayed when the recovering cells were treated with cPDS. Importantly, depletion of cellular ATP levels by inhibition of oxidative phosphorylation or glycolysis (2DG and CCCP treatments, respectively) impeded rG4 unfolding upon starvation removal in a manner similar to the treatment with G4 stabilizer cPDS during recovery phase (**Fig. 4d**), suggesting that the dynamic unfolding of stress-induced rG4s is an ATP-dependent process.

## Discussion

While the formation of rG4s *in vitro* is a well-studied phenomenon, the regulation and function of rG4 folding in living eukayotic cells is not well understood. Our data suggests that while rG4s are mostly unfolded under normal conditions, their folding can be rapidly induced by stress. Based on the evidence that RNA structural elements could play a significant role in RNA stability^31^, protein recognition^32^, and RNA-protein granule assembly (e.g., stress granules and P-bodies)^33-35^, stress-induced rG4 folding could play direct functional roles in human cells. We demonstrated a stress dependent shift in the overall rG4 landscape with an increased level of rG4 formation under a variety of stress conditions (**Fig. 1** and **figs. S1-2**). Our RT stalling analysis on DMS modified total cellular mRNA identified individual folded rG4s in the cells. Of particular interest is the enrichment of stress responsive rG4 elements in 3’UTRs. The rapid folding of such rG4s under stress modulates the stability of specific mRNAs and tested mRNAs show significantly longer half-lives in stressed cells suggesting the contribution of 3’UTR rG4s in mRNA stability during stress. We hypothesize that such global rG4 folding aims on the preservation of rG4-bearing transcripts during stress.

Although cells respond to environmental stress in diverse ways, such stress-induced changes are reversible, and inability to revert gene expression to homeostasis upon resolution of stress contributes to disease^36^. Consistent with this central tenet, we showed that stress induced G4 assembly is reversible upon stress removal, implicating G4 folding as a component of an adaptive stress response program. We propose a model (**Fig. 4e**) that proposes that the reversible stress mediated increase in rG4s folding contributes to selective mRNA stability. Our results also show that rG4 unfolding upon stress resolution depends on the cellular energy status suggesting that rG4 disassembly is an active rather than passive process that relies on the action of ATP-dependent machinery, likely RNA helicases. In summary, we demonstrated a bona fide presence of rG4s in the 3’-UTRs of mRNAs and demonstrated their role as a positive regulator of mRNA stability under cellular stress.

## Acknowledgments

We thank members of Ivanov and Anderson labs for valuable discussion.

Dr. Anderson serves on the Scientific Advisory Boards of Simcere Pharmaceutical and Sedec Therapeutics.

## Author contributions

Conceptualization: PI, PJA, JUG

Methodology: PK, MF, EMV, AVK, JUG

Investigation: PK, MF, EMV, AVK, JG

Funding acquisition: PI, PJA, JUG

Project administration: PI, JUG

Supervision: PI, PJA, JUG, AVK

Writing – original draft: PI, PK, AVK

Writing – review & editing: PI, PJA, JG, PK

## Data and materials availability

All data are available in the main text or the supplementary materials

## Supplementary Materials

Materials and Methods

Supplementary Text

Figs. S1 to S6

Tables S1 to S2

## Supplementary materials for

### Materials and methods

#### Cell culture, cellular stress and *in vivo* DMS treatment

U2OS cells were maintained at 37°C in a CO_2_ incubator in DMEM (Corning) supplemented with 10% FBS, 20 mM HEPES (Gibco), 1% penicillin/streptomycin. To induce the oxidative stress, cells were treated with 100 uM sodium arsenite (Ars-) in regular media for an hour. Starvation stress was induced by incubating the cells in HBSS media with Ca^++^ and Mg^++^ for 2h.

Dimethylsulfate (Sigma) was mixed with ethanol (1:1) and was added to U2OS cells cultured in 15 cm dishes to a final DMS concentration of 8%, and evenly distributed by slow swirling. The cells were incubated for 5 min, the media and excess DMS were decanted, and cells were washed twice with 10 mL of 25% β-mercaptoethanol (Sigma-Aldrich) in PBS to quench any residual DMS. After washing, cells were lysed in 8 mL TRIzol reagent (Invitrogen) supplemented with 5% β-mercaptoethanol, and lysates were immediately processed for mRNA isolation.

#### Immunofluorescence

1×10^5^ U2OS cells were plated into a 24-well plate seed with coverslips. The following day cells were treated as indicated in figure legends. Then the cells were fixed with 4% paraformaldehyde for 15 min, permeabilized with 0.5% Triton-X 100 for 10 min and blocked for 45 min with 5% normal horse serum (NHS; ThermoFisher) diluted in PBS. Primary antibodies were diluted in blocking solution and incubated for 1 h at room temperature or overnight at 4°C. Anti G-quadruplex antibody (BG4) was purchased from Santa Cruz and used at a 1:500 dilution. The cells were washed thrice with PBS and incubated in 1:1000 diluted anti-His tag antibody. Next, cells were washed three times then incubated with secondary antibodies and Hoechst 33258 (Sigma-Aldrich) for 1 h at room temperature and again washed. Coverslips were mounted on glass slides with Vinol and imaged.

#### Microscopy

Wide-field fluorescence microscopy was performed using an Eclipse E800 microscope (Nikon) equipped with epifluorescence optics and a digital camera (Spot Pursuit USB) or an AXIO observer A1 (Zeiss) equipped with epifluorescence optics and a digital camera (SPOT Idea 5.0mp). Image acquisition was done with a 40X objective. Images were merged using Adobe Photoshop and fluorescence intensity was quantified using ImageJ.

Poly(A)-selected RNA or i*n vitro* synthesized RNA oligos (Integrated DNA technologies) in 150 mM KCl or LiCl in T_10_E_0.1_ buffer (10 mM Tris-HCl pH 7.4 and 0.1 mM EDTA) were heated to 95°C for 5 min and then slowly cooled to room temperature.

#### RT-stop profiling and analysis of sequencing reads

DMS-seq and data analysis were performed as previously described (ref 22). Briefly, control or DMS-treated polyA^+^ mRNAs were fragmented by using a buffered zinc solution (Ambion). mRNA fragments were radiolabled by using T4 polynucleotide kinase (NEB) and separated by denaturing PAGE. 60-80 nt fragments were extracted and ligated to a pre-adenylated 3′ adapter containing the RT priming sequence. After non-ligated adapters being removed by denaturing PAGE, RT was performed by using an RT primer containing the 5′ adapter sequence and SuperScript III (Invitrogen) with either 150 mM K^+^ or Li^+^ at 42°C for 10 min. After alkaline hydrolysis of the RNA templates, cDNAs were circularized by using circLigase (Lucigen) at 70°C for 1 hour. Sequencing libraries were generated by PCR and gel-purified. Single-end DNA sequencing was performed on a HiSeq2000 using a standard protocol. Sequencing reads were first de-duplicated using SAMtools. After the molecular barcodes and adapter sequences were trimmed, reads were mapped to a hg19 transciptome reference by Bowtie, requiring zero mismatch and unique mapping. The nucleotide immediately downstream of the most 3′ cDNA nucleotide was assigned as the RT stop.

#### Electrophoretic mobility gel shift assay

RNA oligonucleotides were purchased from Integrated DNA Technologies (IDT). To equilibrate, RNAs were diluted to 10 μM in indicated salt solution, heated to 95 °C for 5 min and slowly allowed to cool to room temperature. For analysis on acrylamide gels, 10 pmol or 50 pmol were ran through a gel and stained with SYBR gold or NMM derivatives, respectively. For SYBR gold staining, gels were post-stained in a solution of 1X SYBR Gold (ThermoFisher Scientific) in 0.5X TBE for 10 min. For NMM staining, gels were post-stained in 100 µM solution of NMM in 0.5X TBE for 30 min. Following staining, gels, were destained for 20 min in 0.5X TBE while rocking at room temperature. Gels were visualized using a 265 nm UV transilluminator.

#### bNMM pull-down protocol

The bNMM G4 pull down experiments were performed in 100 μL volume as follows: bNMM (0, 10 and 50 μM) was mixed with pre-folded fluorescently labelled (3’-FAM) synthetic oligonucleotides (2 μM) in K+ G4 buffer for 1h at room temperature. 20 µL (per pull-down) of streptavidin agarose beads (Pierce) were washed twice with K+ G4 buffer and mixed with 100 µL of rG4 solution. The mixture was tumbled at room temperature for 1 h. Next the bead-RNA mixture was loaded into a Biorad column, washed with 3*1 mL of K+ G4 buffer and NMM bound RNA was extracted with TRIZOL and quantified based upon the fluorescence.

For the cellular mRNA, similar proctocol was adapted except that there was no label in the RNA and the working volume was 750 µL.

#### cDNA synthesis, qPCR, and PCR

mRNA obtained from bNMM pull down experiments was reverse transcribed with the Superscirpt IV first-strand synthesis kit for RT-qPCR (ThermoFisher). The qRT-PCR was performed by using iQ(tm) SYBR® Green Supermix (Bio-Rad), cDNA template, and gene-specific primer sets designed by IDT primer design tool. No reverse transcriptase and no template controls were performed in parallel to check for DNA contamination and primer-dimer. The primer sets used for the study are given in Table 2. Threshold cycle (Ct) values in qRT-PCR experiments were averaged across three biological replicates. The averaged Ct value was used to estimate the enrichment of certain transcripts in the pull-down experiments.

#### 4-thiouracil doping and mRNA decay analysis

U2OS cells were treated with 700 µM 4-thiouracil (4TU) in growth media for different time points. Total RNA was isolated from the cells. Total RNA was divided in two fractions. One fraction of total RNA (7 µg) was reacted with MTSEA-biotin-XXX (Biotium). Biotinylated total RNA was then subjected to streptavidin bead selection. 40 μL streptavidin-agarose (ThermoFisher) were washed with 500 μL of biotin binding buffer (0.1 M NaCl in 10 mM TrisHCl pH 7.4, 10 mM EDTA). The beads were then resuspended in 500 μL binding buffer and mixed with blocking yeast tRNA. Beads were then incubated with gentle agitation for 2h at 4C, washed with 750 μL binding buffer and resuspended in 250 μL biotin RNA binding buffer. MTSEA-biotinylated RNA was then added to the blocked streptavidin beads and incubated with gentle agitation for 4h at 4C. The flowthrough and washes were pooled, sec-butanol concentrated, ethanol-precipitated, washed, and collected as non-biotinylated total RNA, that was used to analyze mRNA decay. 4TU doping, RNA isolation and biotinylation was done in a reduced light environment. Relative RNA was calculated based on the CT value difference between total RNA and non-biotinylated RNA.

#### Starvation rescue experiment

U2OS cells under starvation (as described above) were fed with normal culture medium for different time points and were analyzed for the change in BG4 fluorescence intensity over time.

#### Synthesis and characterization of biotinylated NMM

Common reagents and solvents were purchased from commercial suppliers and used without further purification unless otherwise stated. Tetrahydrofuran (THF) was distilled from sodium-benzophenone under an argon atmosphere. Reaction progress was monitored using analytical thin-layer chromatography (TLC) on pre-coated silica gel GF254 plates (Macherey Nagel GmbH & Co. KG, Düren, Germany), and spots were detected under UV light (254 and 366 nm). Compounds were purified with flash column chromatography with a silica gel and particle size of 40–63 μM (Merck, Darmstadt, Germany) as the stationary phase and hexane/ethyl acetate or dichloromethane/methanol mixtures as eluent systems. Nuclear magnetic resonance spectra were measured on a Bruker AV-400 NMR instrument (Bruker, Karlsruhe, Germany) in deuterated solvents (DMSO-d6 or CDCl3). Chemical shifts are expressed in ppm relative to DMSO-d6 or CDCl3 (2.50/7.26 for 1H; 39.52/77.16 for 13C). The following abbreviations are used to set multiplicities: s = singlet, d = doublet, t = triplet, q = quartet, m = multiplet, br. = broad. Measurements for verification and purity of the compounds were performed by LC/MS. LC–MS/MS data were obtained using a Dionex Ultimate 3000 liquid chromatograph (Dionex, USA) connected to an AB Sciex Qtrap 3200 mass spectrometer (AB Sciex, Canada). LC separation was carried out on a Shim-pack GIST C18-AQ (150 mm × 2.1 mm, 3 µm, Shimadzu, Japan) column. Mobile phase consisted of the mixture of 0.1% (v/v) formic acid in water (A) and acetonitrile (B). Separation was performed in isocratic mode 10% : 90% (A : B). The mobile phase flow rate was 0.3 mL/min. The injection volume was 10 μL. Compounds were detected at λ = 254 nm. All high-resolution mass spectra (HRMS) were measured on AB Sciex TripleTOF 5600+ instrument equipped with DuoSpray (ESI) ion source. Samples were directly injected in the ion source in acetonitrile or methanol solutions acidified by formic acid. The melting points were measured in open capillaries and presented without correction. Synthetic details are available in supplementary materials.

#### Statistical analyses

If not stated otherwise, results are mean values ± standard error of mean (SEM) of at least three independent experiments, or results show one representative experiment of a minimum of three biological replicates. Statistical analyses were performed on all available data. Unless otherwise mentioned, statistical significance was determined using the two-tailed Student’s t test with p values ≤ 0.05 considered statistically significant.

### Synthesis of Biotin-linker (^1^)

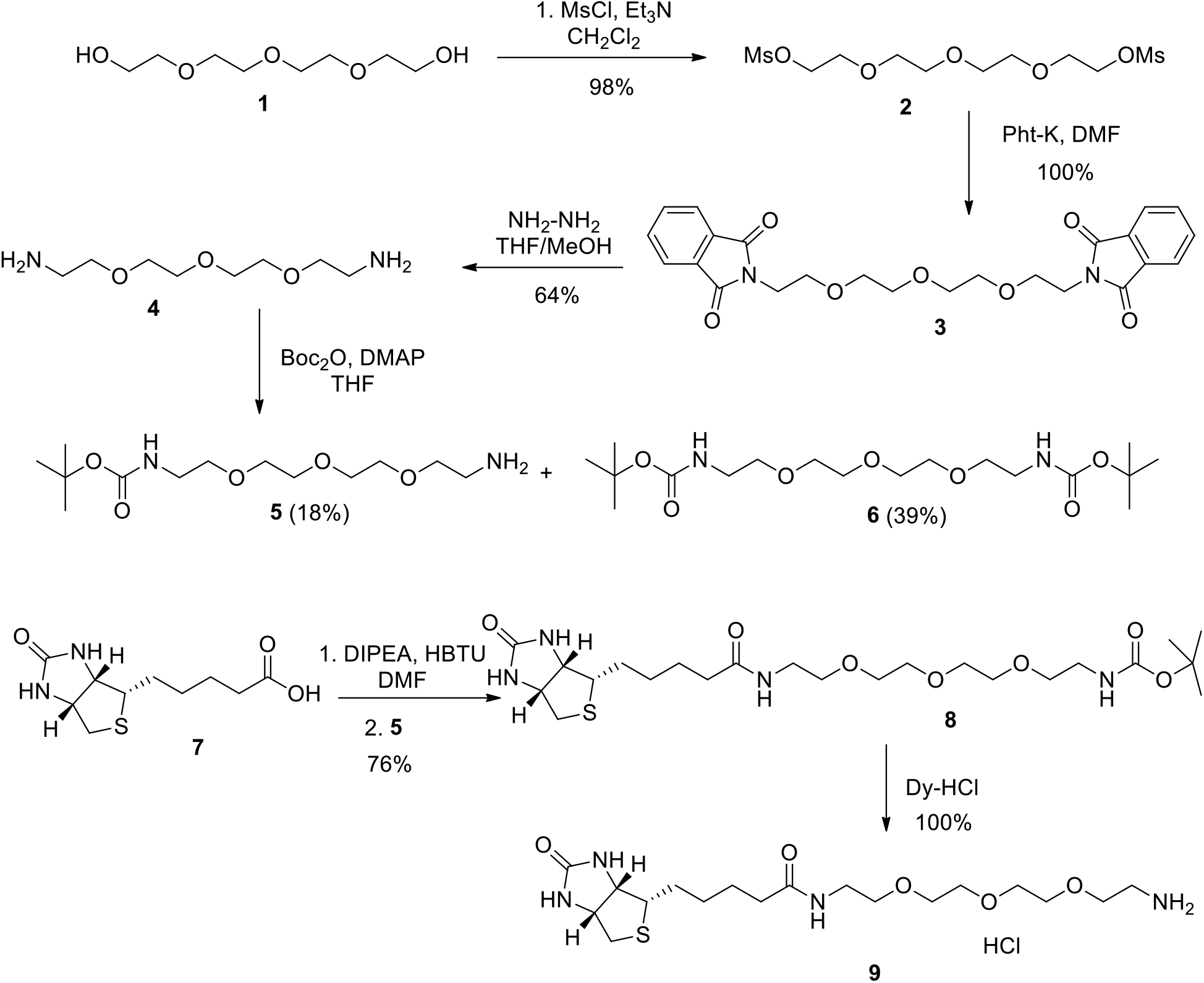

### Tetraethylene glycol dimethanesulfonate (2)

Methanesulfonyl chloride (6 mL, 76.82 mmol) was added dropwise to a stirred solution of PEG4-di-OH **1** (4.455g, 22.59 mmol) and TEA (15 mL, 107.08 mmol) in dichloromethane (20 mL) at 0 ^0^C. After the addition was complete, the resulting mixture was stirred at room temperature for one day. Water was added to quench the reaction. The organic phase was separated, and the aqueous phase was extracted with dichloromethane (2 × 50 mL). The combined organic layers were washed with brine (3 × 60 mL), dried over Na_2_SO_4_, filtered, and the solvent removed under reduced pressure to afford the desired product. Yield 7.75 g (98%).

^1^H NMR (CDCl_3_, 400 MHz) *δ*: 4.37-4.34 (m, 4 H), 3.76-3.73 (m, 4 H), 3.67 -3.60 (m, 8 H), 3.05 (s, 6 H).

### 1,11-Diphthalimido-3,6,9-trioxaundecane (3)

A suspension of 3.03 g (8.65 mmol) **2** and 3.85 g (20.77 mmol) potassium phthalimide in 100 mL abs. DMF was heated for16 h at 80°C. The solvent was removed in vacuum and the residue was suspended in 150 mL DCM and filtrated. The filtrate was washed with water (3 × 100 mL), dried over Na_2_SO_4_, filtrated and concentrated in vacuum. Yield 3.93 g (100%).

^1^H NMR (CDCl_3_, 400 MHz) *δ*: δ = 7.83 (m, 4H, 2 × 5H,6H-Isoindol), 7.69 (m, 4H, 2 × 4H,7H-Isoindol), 3.86 (t, 4H, *J* = 5.7 Hz, 2 x -CH2-CH2-NPht), 3.69 (t, 4H, *J* = 5.7 Hz, 2 x -CH2-CH2-NPth), 3.54 (m, 8H, PhtN-EtO-[(CH2)2-O-]2-Et-NPhth).

### 1,11-Diamino-3,6,9-trioxaundecane (4)

The product **3** (3.0 g, 6.6 mmol) obtained above was dissolved in anhydrous THF–MeOH (1 : 1 (v/v)). To the THF–MeOH solution, 4.0 mL of hydrazine monohydrate were added and the resultant mixture was stirred at room temperature. After 12 h, the white precipitate was removed by filtration. This filtrating process was repeated at least three times to remove the precipitate. The filtrate was concentrated under reduced pressure to give **4** as yellow oil. Yield 0.815 g (164%) (815 mg, 64%).

^1^H NMR (CDCl_3_, 400 MHz) *δ*: 3.61–3.66 (m, 16H).

### Boc-amine-11-Amino-3,6,9-trioxaundecane (5)

The diamine **4** (3.83 g, 0.02 mol, 1 equiv) in DCM solution (10 mL / mmol) was treated with Boc_2_O in default (4.80 g, 0.022 mol, 1.1 equiv) and DMAP (0,024g, 0,2 mmol, 0.01 equiv) for 0.5h at 0 °C. The reaction was stirred at room temperature for 12h. The solvent was removed under reduced pressure. The crude product was purified by silica gel chromatography (EtOAc: hexane 1:1) to yield **5** as a colorless oil (1.08 g, 18% yield) and **6** as a colorless oil (3.04 g, 39% yield).

^1^H NMR (CDCl_3_, 400 MHz) *δ*: 5.27 (s, 1H), 3.65 (m, 12H), 3.52 (q, 3J = 5.9 Hz, 2H), 3.31 (q, 3J = 6.8 Hz, 2H), 2.87 (t, 3J = 5.2 Hz, 2H), 1.45 (s, 9H)

### N-Boc-N’-biotinyl-3,6,9-trioxaundecane-1,11-diamine (8)

Biotin **7** (0.731 g, 2.99 mmol, Sigma, 1.0 equiv) was weighed directly into the reaction flask. This was dissolved in anhydrous DMF (25 mL). HBTU (98%, 0.18 g, 9.00 mmol, 1.1 equiv) was added next. The reaction flask was placed under an argon atmosphere and was stirred for 2 h at room temperature. To this solution was added a solution of **5** (1.05 g, 3.591 mmol, 1.2 equiv) and DIPEA (98%, 0.467 g, 0.61 mL, 0.528 mmol, 1.2 equiv) in anhydrous DMF (10 mL) and the solution mixture was stirred overnight at room temperature under argon to ensure a complete reaction. The reaction mixture was then evaporated and subjected to silica gel chromatography in 9:1 CHCl_3_/MeOH as eluent to afford **6** (1.18 g, 76% yield) as a white solid.

^1^H NMR (CDCl_3_, 400 MHz) *δ*: 6.92 (br t, 1H,CH_2_NHCO), 6.82 (s, 1H, NHbiotin), 6.22 (s, 1H, NH_biotin_), 5.15 (br t, 1H, NH_carbamate_), 4.33 (m, 1 H, CHN H_biotin_), 4.14 (m, 1H, CHN H_biotin_), 3.35 - 3.47 (m, 12H, (CH_2_–O)_6_), 3.26 (br q, 2H, *J* = 4.7, 9.5 Hz, CHS), 3.14 (app q, 2H, *J* = 5.0 Hz, CH_2_NHCO), 2.98 (br q, 1H, *J* = 4.5 Hz, CHS), 2.73 (dd, 1H, *J* = 5.0,13.2 Hz, CHS), 2.58 (d, 1H,*J* = 12.7 Hz, CHS), 2.06 (t, 2H, *J* = 7.4 Hz, CH_2_CONH), 1.51 (m, 4H, (CH_2_)_2biotin_), 1.27 (br s, 11H, (CH_3_)_3_,CH_2biotin_).

### N-Biotinyl-3,6,9-trioxaundecane-1,11-diamine (9)

To a warm solution of compound **8** (0.655 g, 1.26 mmol) in dioxane (4 ml) was added a 4M solution of HCl in dioxane (2 mL). The mixture was stirred at 50 °C for 0.5 h while oil dropped out in the mixture. TLC (10% MeOH in CH_2_Cl_2_) indicated the reaction was complete. The oil was dissolved in methanol and the resulting solution was evaporated, the residue was co-evaporated several times with CHCl_3_, Et_2_O and CH_2_Cl_2_. Yield 0.574 g (100%).

^1^H NMR (DMSO-d_6_, 400 MHz) *δ*: 7.98 - 8.26 (m, 3H), 7.83 - 7.98 (m, 1H), 4.24 - 4.39 (m, 1H), 4.06 - 4.19 (m, 1H), 3.61 (t, *J*=5.2 Hz, 2H), 3.53 (d, *J*=16.3 Hz, 8H), 3.32 - 3.45 (m, 2H), 3.13 - 3.24 (m, 2H), 3.05 - 3.13 (m, 1H), 2.87 - 3.01 (m, 2H), 2.82 (dd, *J*=12.4, 4.98 Hz, 1H), 2.58 (d, *J*=12.5 Hz, 1H), 2.07 (t, *J*=7.3 Hz, 2H), 1.56 - 1.69 (m, 1H), 1.39 - 1.55 (m, 3H), 1.16 - 1.38 (m, 2H).

### Synthesis of NMM-IX

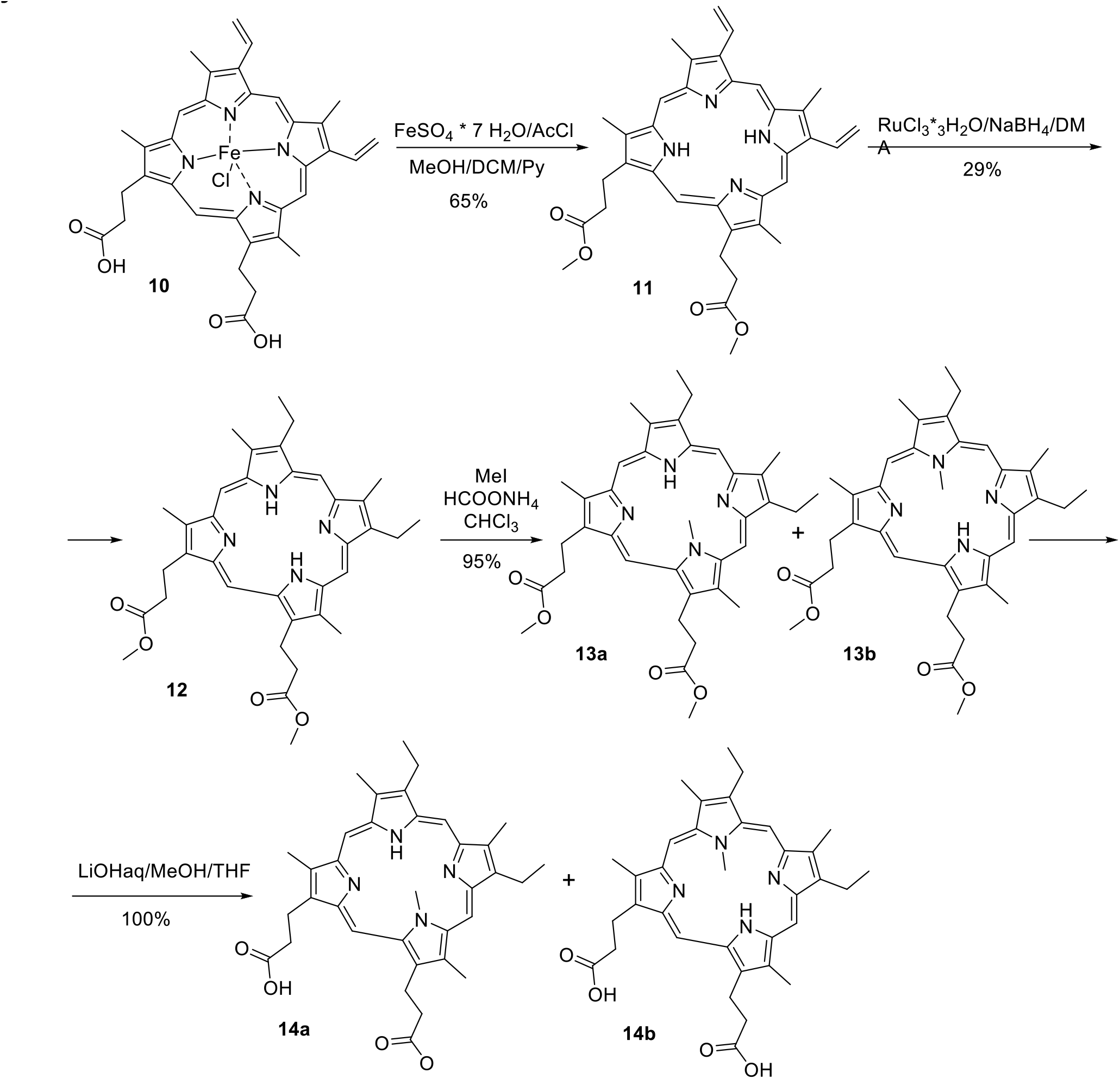

### Protoporphyrin IX dimethyl ester (11)^2^

Hemin (**10**) (0.5 g, 0.76 mmol) and pyridine (0.5 mL) were placed in a three-necked flask, then MeOH (30 ml), CH_2_Cl_2_ (30 ml) and Mohr’s salt (1.42 g, 5.1 mmol) were added. Acetylchloride (15 ml, 21.0 mmol) was gradually added under stirring and cooling, while the temperature was kept below 35 °C. The mixture was stirred for 1 h and then diluted with H_2_O (50 ml). The bottom organic layer was separated, washed with aqueous ammonia (25%, 30 ml), then with H_2_O (20 ml) and dried over anhydrous Na_2_SO_4_. The product was purified by chromatography on silica gel 60 mesh using CH_2_Cl_2_ as the eluent. M.p.: 238 - 240 °C. Yield 0.29 g (65%).

^1^H NMR (CDCl_3_, 400 MHz) *δ*: 9.67 - 9.98 (m, 4 H), 7.99 - 8.25 (m, 2 H), 6.29 (dd, *J*=17.6, 12.32 Hz, 2 H), 6.14 (t, *J*=10.3 Hz, 2 H), 4.22 - 4.39 (m, 4 H), 3.69 (s, 6 H), 3.40 - 3.60 (m, 12 H), 3.13 - 3.32 (m, 4 H)

### Mesoporphyrin IX dimethyl ester (12)^3^

A 100 mL three-neck-round-bottom flask was fitted with a reflux condenser and an oil-bubbler was charged with a solution of **11** (0.591 g, 1.00 mmol, 1 equiv) in DMA (20 mL, 0.05M), RuCl_3_·3H_2_O (0.131 g, 0.50 mmol, 0.5 equiv), then to this mixture NaBH_4_ (0.393 g, 10.40 mmol, 10.40 equiv) was added with portions for three times in less than 30 min under N_2_ atmosphere and then stirred at 25 °C. After 30 min, the UV–vis spectrum of a sample aliquot showed complete conversion. The mixture was flushed with Ar for 5 min, opened to the atmosphere, and concentrated to 5 mL. The result mixture was transformed into 200 mL water and extracted with 300 mL CH_2_Cl_2_. The organic phase was collected and washed for three times with water to remove DMA and Ru salts. The crude product was purified by column chromatography (60-80 mesh silica gel). The filtrate was collected and evaporated to dryness to afford **12** as brownish, red or black solids. M.p.: 211 - 213 °C. Yield: 0.25 g (42%).

^1^H NMR (400 MHz, CDCl_3_) *δ*: 10.21 (s, 1H, 20-H), 10.10 (s, 2H, 5-, 15-H), 9.93 (s, 1H, 10-H), 4.29 - 4.58 (m, 4H, 13-,17-*β*-CH2, *α* to carbonyl), 4.12 (q, *J*_HH_=7.6 Hz, 4H, 3-, 8-CH_2_CH_3_), 3.54 – 3.82 (M, 18H, 2-, 7-, 12-,18-CH_3_ and 13-, 17-OCH_3_), 3.29 (t, *J*_HH_=7.5 Hz, 4H, 13-, 17-*α*-CH2, *α* to carbonyl), 1.89 (t, *J*_HH_=7.6 Hz, 6H, 3-, 8-CH2CH3), 3.71 (s, 2H, NH).

### N-Methylmesoporphyrin IX dimethyl ester (13)^4^

Mesoporphyrin IX dimethyl ester **12** (0.242 g, 0.407 mmol, 1.0 equiv) was dissolved in CHCl_3_, (20 mL, 0.02M) and MeI (3.9 mL, 8.886 g, 62.6 mmol, 154.0 equiv) and excess ammonium formate (0.256 g, 4.065 mmol, 10.0 equiv) was added. The mixture was boiled under reflux for 12 hr. cooled and the inorganic salts removed by filtration. The filtrate was concentrated and purified by column chromatography (60-80 mesh silica gel). MeOH-DCM (1:10) eluted a red-brown fraction which on work up and crystallization from CHCl_3_-MeOH afforded N-Methylmesoporphyrin IX dimethyl ester (**13**) as the mixture of two products (**13a** and **13b**) as purple plates. M.p.: 245 - 250 °C. Yield: 0.23 g (95%). R_f_ = 0.2 (CH_2_Cl_2_/MeOH 20:1). **13a**. rt = 1.91 min. LC-MS: m/z [M+H]+ = 609 Da, **13b**. rt = 1.98 min. LC-MS: m/z [M+H]+ = 609 Da.

^1^H NMR (400 MHz, CDCl_3_) *δ*: 10.34 - 11.01 (m, 4 H), 3.74 - 4.80 (m, 15 H), 3.50 - 3.73 (m, 13 H), 3.27 - 3.50 (m, 4 H), 2.97 - 3.08 (m, 1 H), 1.92 - 2.02 (m, 3 H), 1.88 (t, *J*=7.6 Hz, 1H), 1.50 (td, *J*=7.6, 2.60 Hz, 2H).

### NMM-IX (14)^5^

N-Methylmesoporphyrin IX dimethyl ester **13** (0.14 g, 0.23 mmol, 1.0 equiv) was dissolved in a THF/MeOH/H_2_O mixture (15 mL, 1:1:1). LiOH·H2O (0.096 g, 2.3 mmol, 10.0 equiv) was added and the reaction mixture was stirred for 8 h at room temperature. The reaction mixture was filtered and the volume of the filtrate was reduced under vacuum (5 mL). Citric acid (1N, 2 mL) was slowly added to this solution with stirring to precipitate the dark violet solid, which was filtered off and then washed with water (5 mL) to give **14** as a dark violet solid (the mixture **14a** and 1**4b**). Yield: 0.134 g (100%). Rf = 0.8 (CH_2_Cl_2_/MeOH 1:1). **14a**. rt = 1.67 min. LC-MS: m/z [M+H]+ = 581 Da, **14b**. rt = 1.68 min. LC-MS: m/z [M+H]+ = 581 Da.

^1^H NMR (400 MHz, CDCl_3_) *δ*: 11.86 - 12.93 (m, 2 H), 9.72 - 10.98 (m, 4 H), 3.40 - 4.83 (m, 19 H), 2.55 - 3.30 (m, 6 H), 1.70 - 1.91 (m, 4 H), 1.18 - 1.34 (m, 2 H)

### Synthesis of biotinylated NMM derivatives

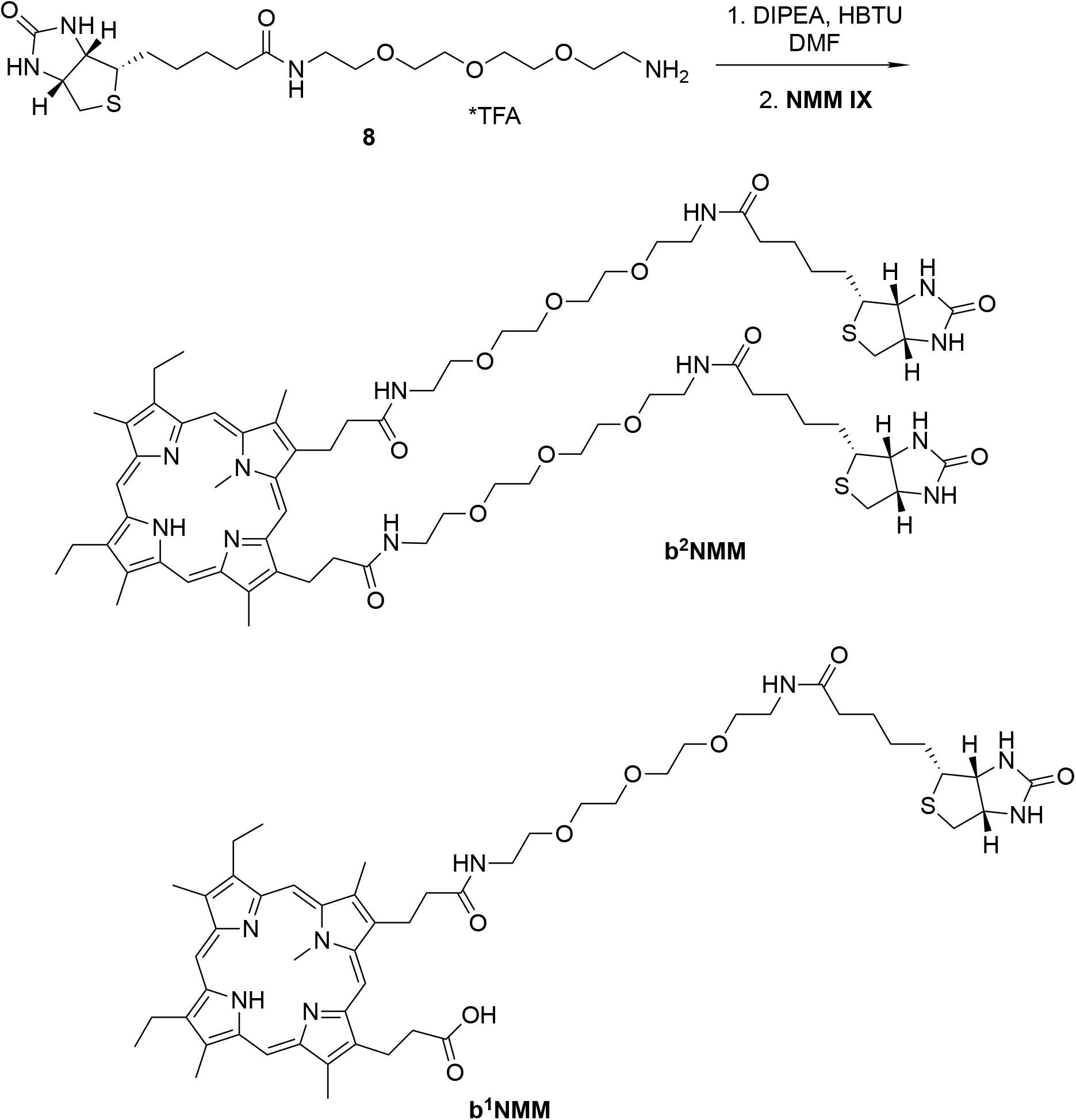

#### Preparation of b^2^NMM and b^1^NMM

HBTU (98%, 0.18 g, 9.00 mmol, 2.2 equiv) was added to a solution of NMM-IX **14** (0.123 g, 0.211 mmol, 1.0 equiv) in anhydrous DMF (5 mL), which was stirred for 2 h at room temperature. To this solution was added a solution of **9**·HCl (0.294 g, 0.561 mmol, 2.66 equiv.) and DIPEA (98%, 0.07 g, 0.09 mL, 0.528 mmol, 2.5 equiv) in anhydrous DMF (4 mL) and the solution mixture was stirred for 48 h at room temperature. After filtering, the filtrate was evaporated to dryness. The residual solid was dissolved in CH_2_Cl_2_ and the solution was washed with water, dried over anhydrous Na_2_SO_4_, and evaporated under reduced pressure to obtain a purple solid. The crude material was purified via preparative LC/MS with the following conditions: Column: YMC-Actus Pro C18, 20×250 mm, 12-μm particles; Mobile Phase A: 2:98 CH_3_CN:H_2_O with 0.1% formic acid; Mobile Phase B: 95:5 CH_3_CN:H_2_O with 0.1% formic acid; Gradient: 25-65% B over 15 minutes, then a 5-minutehold at 100% B; Flow: 20 mL/min. Fractions containing the desired products were combined and dried via centrifugal evaporation.

The yield of the **b^2^NMM** was 35.6 mg, and its estimated purity by LCMS analysis was 100%. Retention time=2.22 min; HRMS (ESI-MS) calcd for C_71_H_105_N_12_O_12_S_2_ [M+H]^+^ 1381.7411, found 1381.7338.

The yield of the **b^1^NMM** was 8.7 mg, and its estimated purity by LCMS analysis was 100%. Retention time=2.31 min; HRMS (ESI-MS) calcd for C_53_H_73_N_8_O_9_S [M+H]^+^ 997.5222, found 997.5216.

**Table SI 1.**
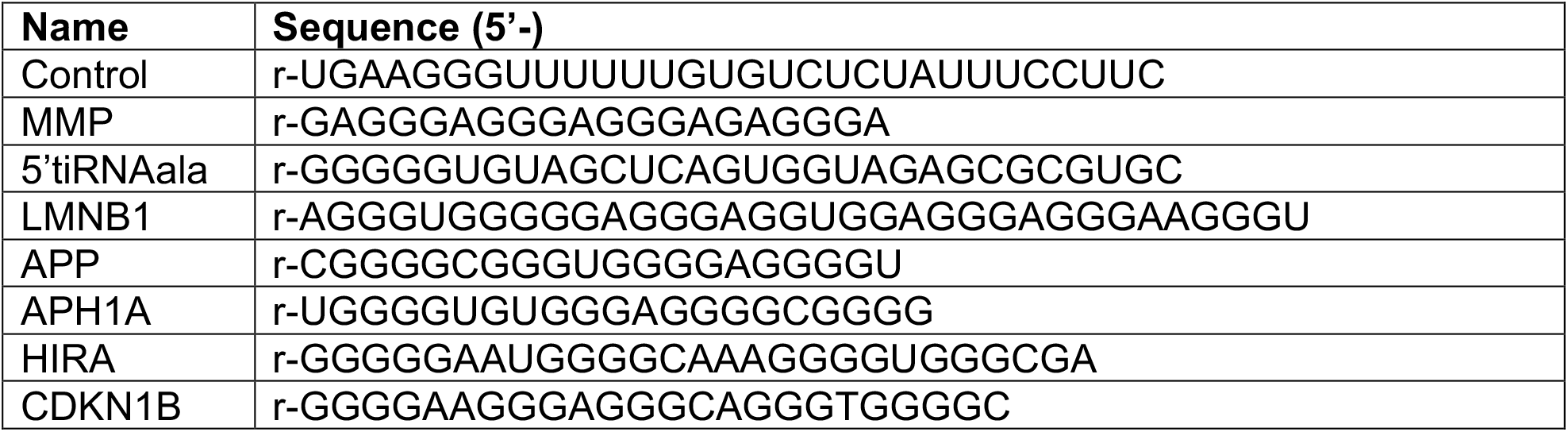
List of the rG4/ non-rG4 oligos.

**Table SI 2.**
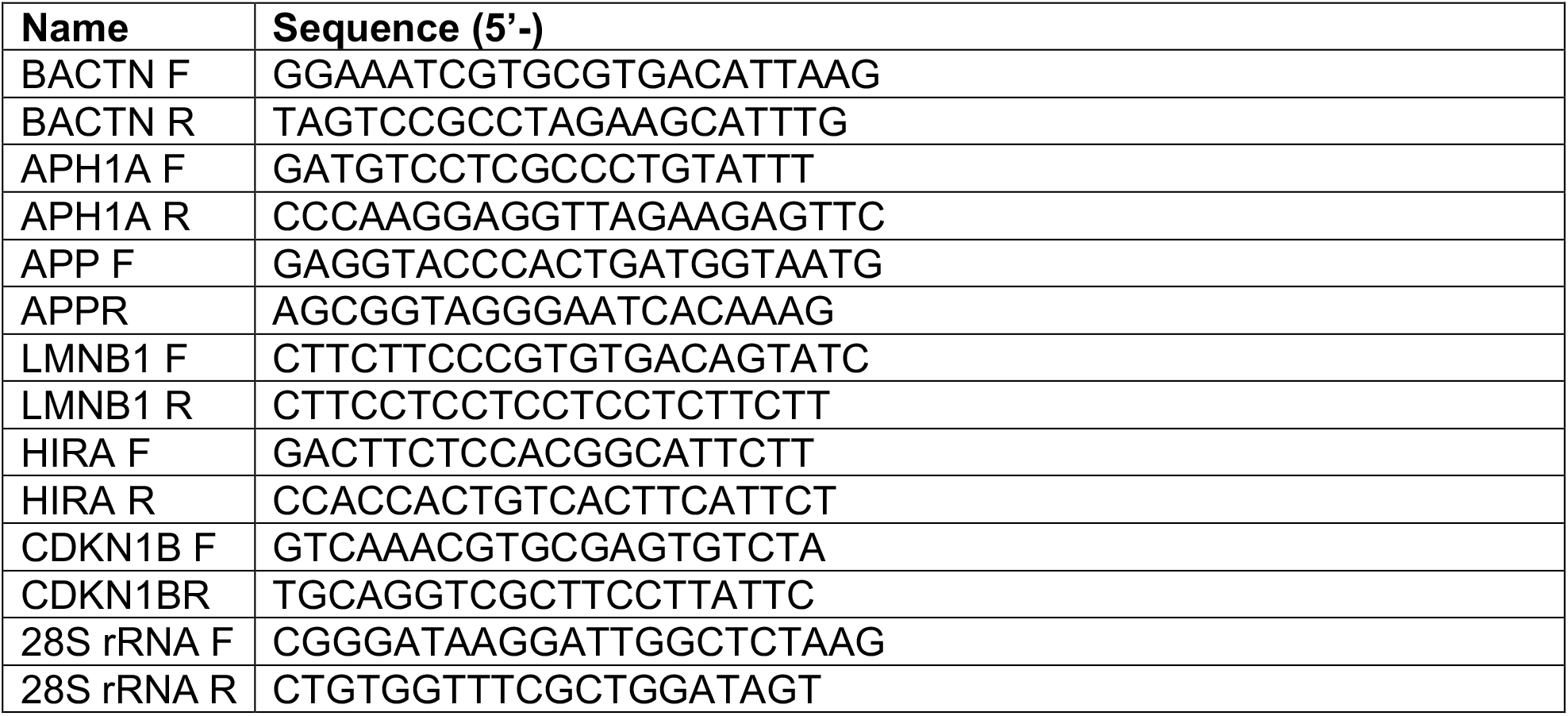
List of the primers.

**Figure S1.**
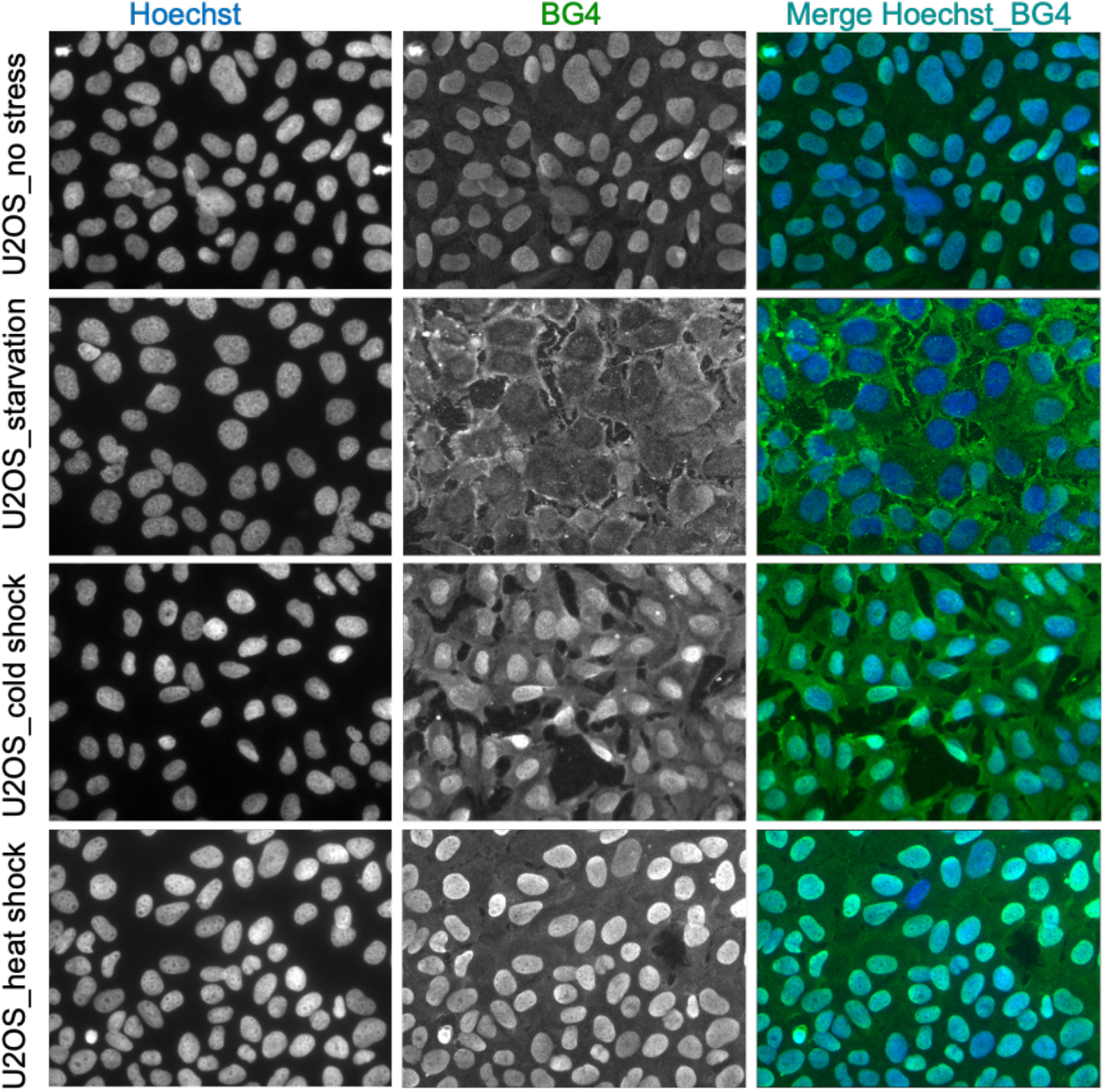
Increased rG4 formation under different cellular stresses U2OS cells.

##### Increased rG4 formation under different stresses in U2OS and COS7 cells

We studied the stress response of rG4 formation using cold-shock and heat shock stresses both in U2OS cells and COS7 cells. Cold shock (10C overnight) as well as starvation stress seems to affect rG4 formation similarly in both cell types. We also observed a slightly enhanced BG4 signals with heat shock (42C for 30 min) in both the cell types.

**Figure S2.**
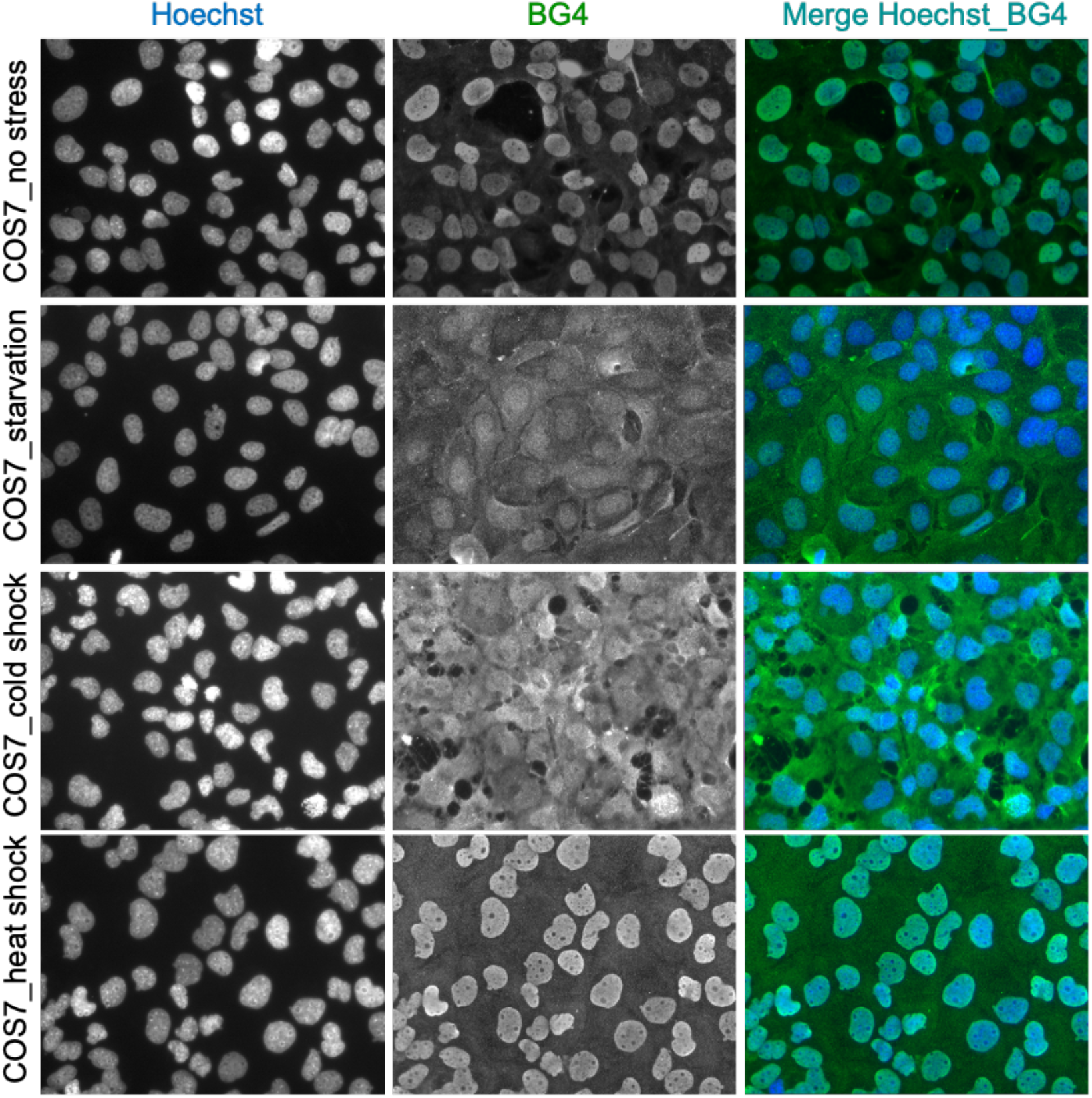
Increased rG4 formation under different cellular stresses in COS7 cells

**Figure S3.**
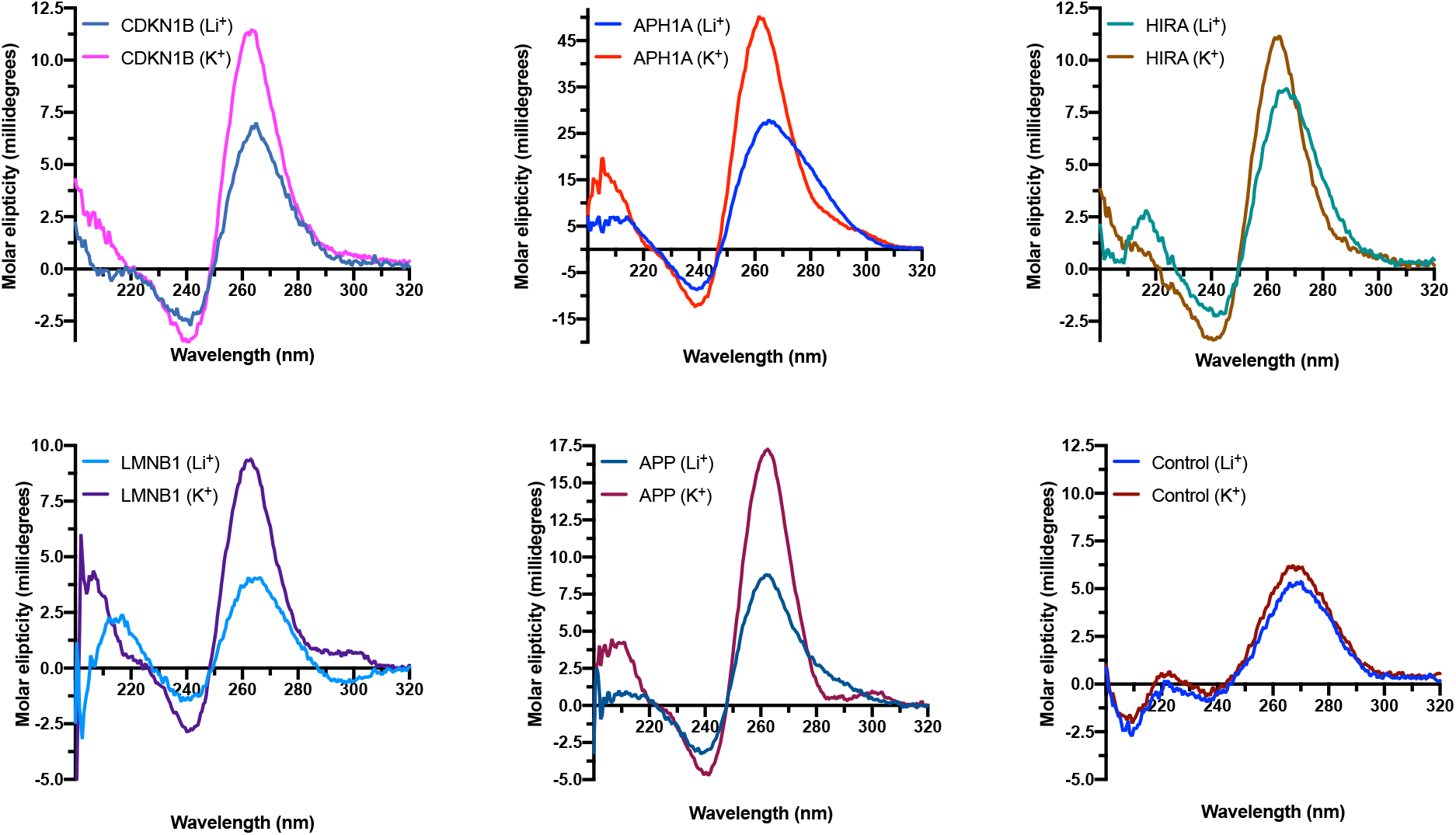
CD spectra of candidate oligos show a rG4 feature which is absent in the control oligo.

##### Circular dichroism (CD) characterization of rG4 formation by DMSeq identified mRNA oligos

JASCO J815 spectropolarimeter was used to collect CD spectra. The oligonucleotides were dissolved in 150 mM K+ in T_10_E_0.1_ buffer. Quartz cuvettes with a 1 mm path length were used with the sample volumes of 200 μL to achieve a sample concentration of 5 μM. Spectra were collected in the range between 220 and 320 nm at 20 °C from three scans and a buffer baseline was subtracted from each spectrum. CD was expressed as the difference in the molar absorption of the right-handed and left-handed circularly polarized light. An increased peak intensity of the oligo under K+ environment at 260-265 nm and a trough at 240 nm, that shows a reduced peak intensity under Li+ environment suggests the formation of a G4. All the selected candidates show the G4 characters in CD while the control oligo that cannot form G4 structure shows no change in the CD behavior.

**Figure S4.**
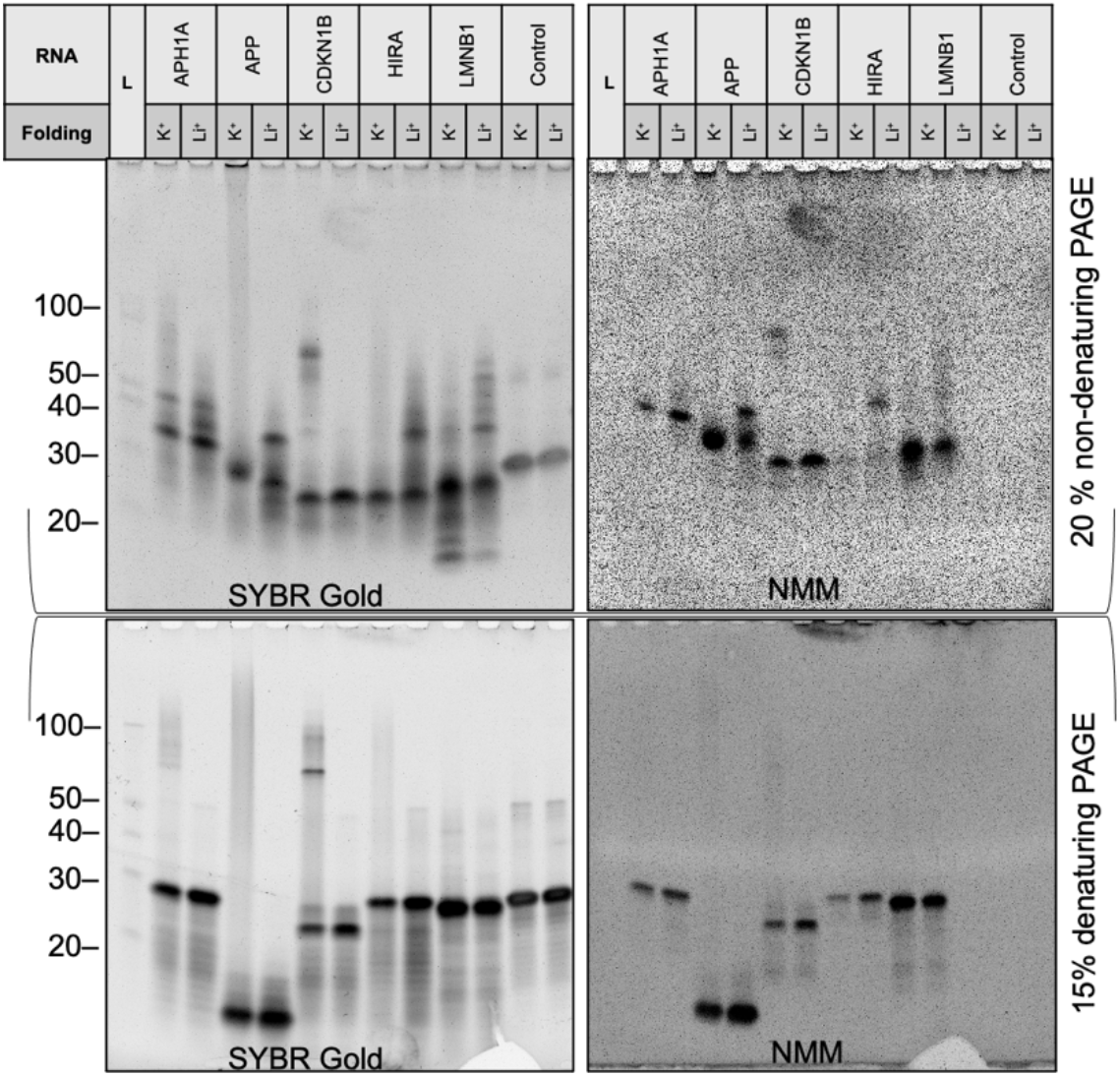
Electromobility gel shift assay coupled with G4 specific dye (N-methyl mesoporphyrin IX, NMM) staining indicate the formation of rG4s by candidate oligos even under harsh conditions (Li^+^ and denaturing PAGE). SYBR Gold detects nucleic acids non-selectively while NMM selectively binds to parallel G4s.

**Figure S5.**
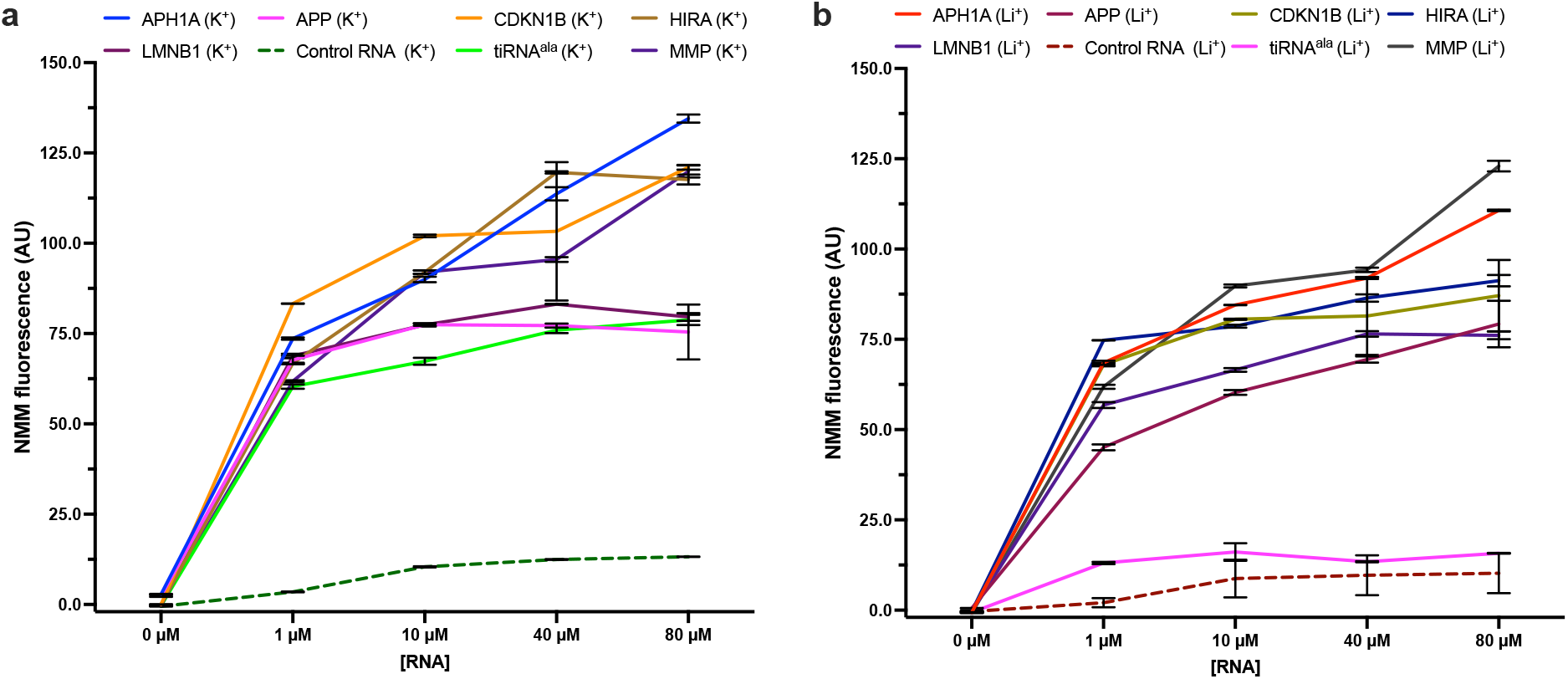
NMM fluorescence assay further confirms the formation of rG4s by candidate oligos even under harsh conditions (Li^+^). Control RNA is a non-rG4 oligo, and 5’tiRNA^ala^ and MMP are positive controls (It has been previously shown that 5’tiRNA^ala^ mostly stays in non-rG4 form under Li^+^ environment while MMP can stay in rG4 form even under Li^+^ environment.

**Figure S6.**
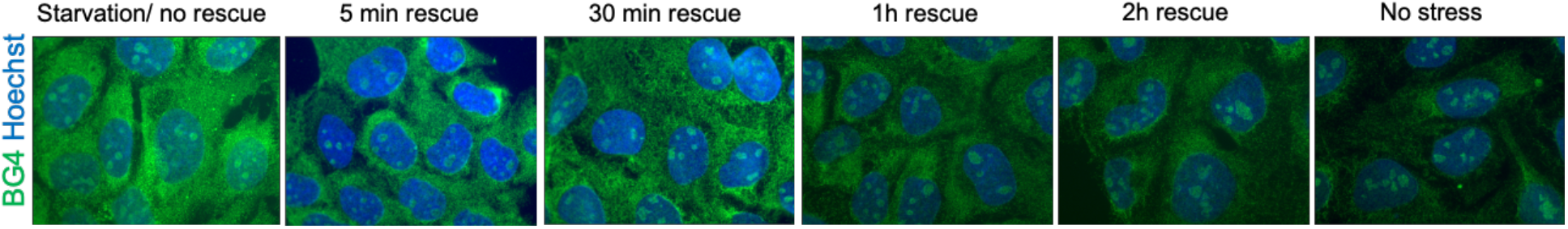
Represented images to illustrate the reversibility of starvation induced rG4 folding upon rescue of the stress (Quantification of the three independent experiments is presented in **Fig. 4c**).

## Footnotes

1 N-Boc-N’-biotinyl-3,6,9-trioxaundecane-1,11-diamine (**8**) was prepared by a method described in: Johanna Davila, Armelle Chassepot, Johan Longo, Fouzia Boulmedais, Andreas Reisch, Benoît Frisch, Florent Meyer, Jean-Claude Voegel, Philippe J. Mésini, Bernard Senger, Marie-Hélène Metz-Boutigue, Joseph Hemmerlé, Philippe Lavalle, Pierre Schaaf and Loïc Jierry, *J. Am. Chem. Soc*, 2012, vol. 134, # 1, p. 83 - 86. dx.doi.org/10.1021/ja208970b

2 Protoporphyrin IX dimethyl ester (**11**) was prepared by a method described in: Natalya Sh. Lebedeva, Elena S. Yurina, Yury A. Gubarev, Aleksey N. Kiselev and Sergey A. Syrbu. Mendeleev Commun., 2020, 30, 211–213. dx.doi.org/10.1016/j.mencom.2020.03.027

3 Mesoporphyrin IX dimethyl ester (**12**) was prepared by a method described in: Júlio S. Rebouças and Brian R. James. Tetrahedron Lett., 2006, 5119–5122. dx.doi.org/10.1016/j.tetlet.2006.05.083

4 N-Methyl mesoporphyrin IX dimethyl ester (**13**) was prepared by a method described in: Ajita M. Abeysekera, Ronald Grigg, Jadwiga Trocha-Grimshaw. Tetrahedron, 1980, 36, 1857-1868

5 NMM-IX (**14**) was prepared by a method described in: Mohamed E. El-Zaria, Hyun Seung Ban, Hiroyuki Nakamura, Chem. Eur. J. 2010, 16, 1543 – 1552. dx.doi.org/10.1002/chem.200901532

